# Investigating the neuroprotective effect of AAV-mediated β-synuclein overexpression in a transgenic model of synucleinopathy

**DOI:** 10.1101/300822

**Authors:** Dorian Sargent, Dominique Bétemps, Matthieu Drouyer, Jérémy Verchere, Damien Gaillard, Jean-Noël Arsac, Latifa Lakhdar, Anna Salvetti, Thierry Baron

## Abstract

Parkinson’s disease (PD) and multiple system atrophy (MSA) are neurodegenerative diseases characterized by inclusions mainly composed of α-synuclein (α-syn) aggregates. The objective of this study was to investigate if β-synuclein (β-syn) overexpression could have beneficial effects by inhibiting the aggregation of α-syn. The M83 transgenic mouse is a model of synucleinopathy, which develops severe motor symptoms associated with aggregation of α-syn. M83 neonate or adult mice were injected with adeno-associated virus vectors carrying the human β-syn gene (AAVβ-syn) or green fluorescent protein gene (AAVGFP) using different injection sites. One or two months later, M83 disease was accelerated or not using brain M83 extracts from mouse (M83) or human (MSA) origins. AAV mediated β-syn overexpression detected by ELISA did not delay the disease onset, regardless of the AAV injection route and of the inoculation of brain extracts. Accordingly, phosphorylated α-syn levels detected by ELISA in sick mice were similar after injecting AAVβ-syn or AAVGFP. Instead, immunohistochemistry analysis of β-syn indicated the presence of proteinase-K resistant β-syn staining specifically in sick M83 mice inoculated with AAVβ-syn. This study indicated for the first time that β-syn could form aggregates in a model of synucleinopathy when it is expressed by a viral vector.

## Introduction

Parkinson’s disease (PD), dementia with Lewy bodies (DLB) and multiple system atrophy (MSA) are synucleinopathies, characterized by inclusions mainly composed of an aggregated form of α-synuclein (α-syn) in the central nervous system (CNS). As for prions, aggregated forms of α-syn propagate within the CNS during the associated neuro-degenerative diseases. This characteristic was initially suggested in humans by Braak’s description of PD stages, and was confirmed later in experimental models of synucleinopathies, in particular in the M83 transgenic mouse model^1^. M83 mice express the human A53T mutated α-syn found in some familial PD forms, under the control of the mouse prion promoter^1^. These mice spontaneously develop severe motor impairment at 8-16 months of age at the homozygous state. The symptomatology is associated with the accumulation in the CNS, of a pathological form of α-syn (α-syn^P^), heavily phosphorylated at serine 129 residue. We previously showed that, in M83 mice, disease onset can be accelerated by the intracerebral inoculation of brain homogenates from sick M83 mice^2^. This model was further characterized by development of an original ELISA that specifically detects, and allows to easily quantify, the pathological α-syn in sick M83 mice^3-5^. This *in vivo* model may be useful for testing novel therapeutic strategies, particularly targeting progression of the α-syn aggregation.

β-synuclein (β-syn) is another member of the synuclein family, lacking a part of the non-amyloid component, a specific region suggested to be amyloidogenic in α-syn. According to *in vitro* studies, unlike α-syn, β-syn alone is not able to form aggregates, but could instead interact with α-syn and reduce its capacity to aggregate^6^. These anti-aggregative features have been investigated i*n vivo*, and some of these studies indeed reported a neuroprotective effect and reduction of α-syn inclusions after overexpression of the human β-syn mediated by DNA microinjection or lentiviral vectors in the transgenic D mouse model, overexpressing human wild-type α-syn^7,8^. Another study also suggested that crossing mice overexpressing human β-syn with M83 mice delayed the M83 disease onset and reduced the α-syn aggregation^9^. However, two recent studies by Taschenberger *et al* and Landeck *et al* which analyzed the impact of human β-syn overexpression mediated by adeno-associated viral vectors (AAV) on nigral dopaminergic neurons in rats, described a neurodegeneration with β- syn aggregates^10,11^.

Here, in order to assess the effects of β-syn overexpression in the context of these recent and unexpected data suggesting that β-syn produced using AAV may be able to aggregate, we investigated the impact of human β-syn overexpression mediated by AAV on the onset of the synucleinopathy of M83 mice. Two strategies of inoculation of the AAV vector were sequentially tested: (i) an intracerebroventricular (ICV) inoculation at birth, in order to generate a widespread overexpression of the vector in the central nervous system^12^, and (ii) an inoculation of the AAV vector into the ventral tegmental area (VTA)^13^, a region of the mesencephalon in which prominent α-syn aggregation occurs in sick M83 mice^1^. Our results indicate that, regardless of the AAV inoculation strategy, continuous AAV-mediated β-syn overexpression in neurons did not delay the onset of the disease and did not modify the α-syn^P^ levels measured by ELISA, strongly suggesting that β-syn may not protect against α-syn aggregation and propagation.

## Results

To overexpress β-syn in the M83 CNS, we used self-complementary (sc)AAV9 vectors expressing human β-syn (AAVβ-syn) gene or, as a control, the green fluorescent protein (AAVGFP) gene under the control of the human synapsin1 promoter and with a post-transcriptional WPRE regulatory sequence (Supplementary figure 1A).

**Figure 1.**
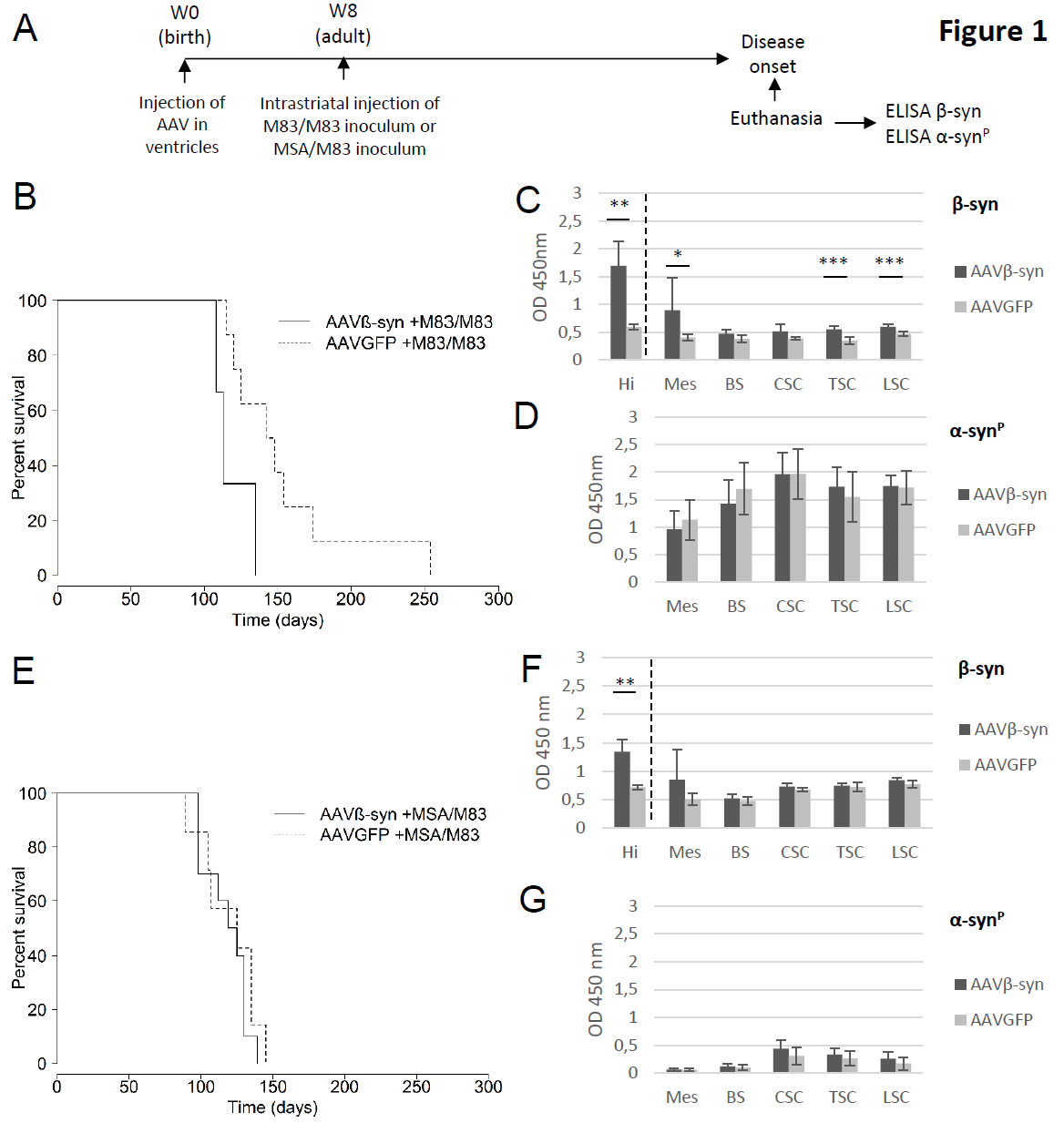
Impact of the intracerebroventricular inoculation of AAV on the accelerated disease of M83 mice. (A) M83 mice were injected at birth (Week 0) with AAVβ-syn or AAVGFP vectors per lateral ventricle and challenged two months later (Week 8) by the injection of M83/M83 (B-D) or MSA/M83 (E-G) inoculum in the striatum to accelerate the disease. After the appearance of the first specific symptoms of M83 disease (from 3 months to 8 months after challenge), mice were sacrificed and dissected in order to perform biochemical analysis. (B) M83 disease-associated survival after the inoculation of the M83/M83 brain extract is represented (significant difference according to log rank test, p<0.05). (C, D) Quantification of total β-syn (mouse β-syn and human β-syn produced by the vector) (C) or of pathological Ser129-phosphorylated form of α-syn (α-syn^P^) (D) by ELISA in CNS regions of the same sick M83 mice groups (n=6 for the treated and the control group). (E) M83 disease-associated survival after the inoculation of the MSA/M83 brain extract (no significant difference according to log-rank test). (F, G) Quantification of total β-syn (F) or α-syn^P^ (G) by ELISA in CNS regions of the same sick M83 mice groups (n=6 for the treated group and n=5 for the control group). Hi: hippocampus, Mes: mesencephalon, BS: brain stem, CSC: cervical spinal cord, TSC: thoracic spinal cord, LSC: lumbar spinal cord. Data are shown as mean ± sd. *p<0,05; **p<0,01; ***p<0,001 according to Student test (B, C) or Wilcoxon test (E, F).

To functionally evaluate the AAV vectors, we first inoculated AAVβ-syn in the right and left cerebral ventricles of wild-type B6C3H newborn mice (genetic background of M83 transgenic mice) to obtain a widespread expression of the transgene in the CNS^12^. One month later, using viral specific primers, vector mRNA was detected in all regions of the brain and in the spinal cord (Supplementary figure 1B), as described^14^, with higher levels in rostral regions of the brain (olfactory bulbs, cortex, hippocampus, striatum) and mesencephalon.

AAV vectors were first tested in M83 mice challenged by inoculation of brain extracts from sick M83 mice, an experimental design which is characterized by a quicker onset of the disease with a lower intragroup variability as compared to unchallenged M83 mice^2^. AAV vectors were ICV inoculated at birth and two months later challenged with different brain extracts of sick M83 mice, one derived from a second passage of a sick M83 brain extract in M83 mice (M83/M83 inoculum), and the other from a second passage of a human brain extract from a MSA patient in M83 mice (MSA/M83 inoculum) (Figure 1A)^5^. M83 mice were euthanized after the detection of the first symptoms of disease, *i.e* balance disorders or hind limb paralysis, identified by two independent observers.

In both experiments, no delay in the onset of the disease was observed in AAVβ-syn injected mice as compared to control AAVGFP mice, regardless of the type of inoculum (M83/M83 or MSA/M83). Instead, AAVβ-syn injection significantly accelerated the onset of the disease in animals challenged with the M83/M83 extract as compared to control mice (log rank test p<0,05) (Figure 1B). In contrast, no difference was observed in animals challenged with the MSA/M83 extract (Figure 1E). After dissection of the brains, biochemical analyses were realized. As the vector expression was higher in the rostral regions of the brain than in other CNS regions after ICV injections, we first checked the vector expression by detecting total β- syn in the hippocampus. An ELISA test adapted from Krassnig *et al.* allowed to detect a significant overexpression of β-syn (both murine β-syn and human β-syn produced by the vector) in the hippocampus, confirming the continuous expression of the AAVβ-syn until the disease onset (Figure 1C, and F). We next focused on the mesencephalon, the brainstem and the spinal cord because these CNS regions are known to be strongly positive in α-syn^P^ in sick M83 mice^3-5^. A significant overexpression of β-syn was only detected in the mesencephalon and in the spinal cord of sick M83 mice injected with AAVβ-syn challenged with M83/M83 inoculum, but not in mice challenged with the MSA/M83 inoculum (Figure 1C and F). Since the M83 disease is associated with moderate or high α-syn^P^ levels into different brain regions and in the spinal cord^5^, we quantified α-syn^P^ using ELISA in order to detect a potential effect of β-syn on this biochemical marker of the disease. Our analyses did not show any significant difference in the α-syn^P^ levels in any of the examined CNS regions after AAVβ-syn inoculation, regardless of the nature of the inoculum used to accelerate the disease (Figure 1D, and G).

To further study the levels of β-syn protein after AAV injections in M83 mice throughout the CNS, we used Western blot and ELISA which allowed to detect total β-syn in specific brain regions. Comparing total β-syn levels of 5 to 7 months old sick M83 mice inoculated at birth with AAVβ-syn or AAV GFP then challenged with the MSA/M83 inoculum (same mice as in Figure 1E), an overexpression of β-syn protein was detected by ELISA in the olfactory bulbs, cortex, striatum, hippocampus, but not in the other regions (Figure 2A), consistent with the results obtained by qRT-PCR at one month after AAV inoculation (Supplementary figure 1B). Western blot analyses confirmed an overexpression of β-syn protein in the hippocampus of the same mice after AAVβ-syn inoculation (Figure 2B and C). Most of the animals inoculated with AAVβ-syn showed a β-syn overexpression detectable until the disease onset (Figure 1C, F). Only a few animals did not show significant overexpression of β-syn detected by ELISA or by immunohistochemistry (5/48 animals, in the entire study); these animals were excluded for the study analyzing the onset of disease.

**Figure 2.**
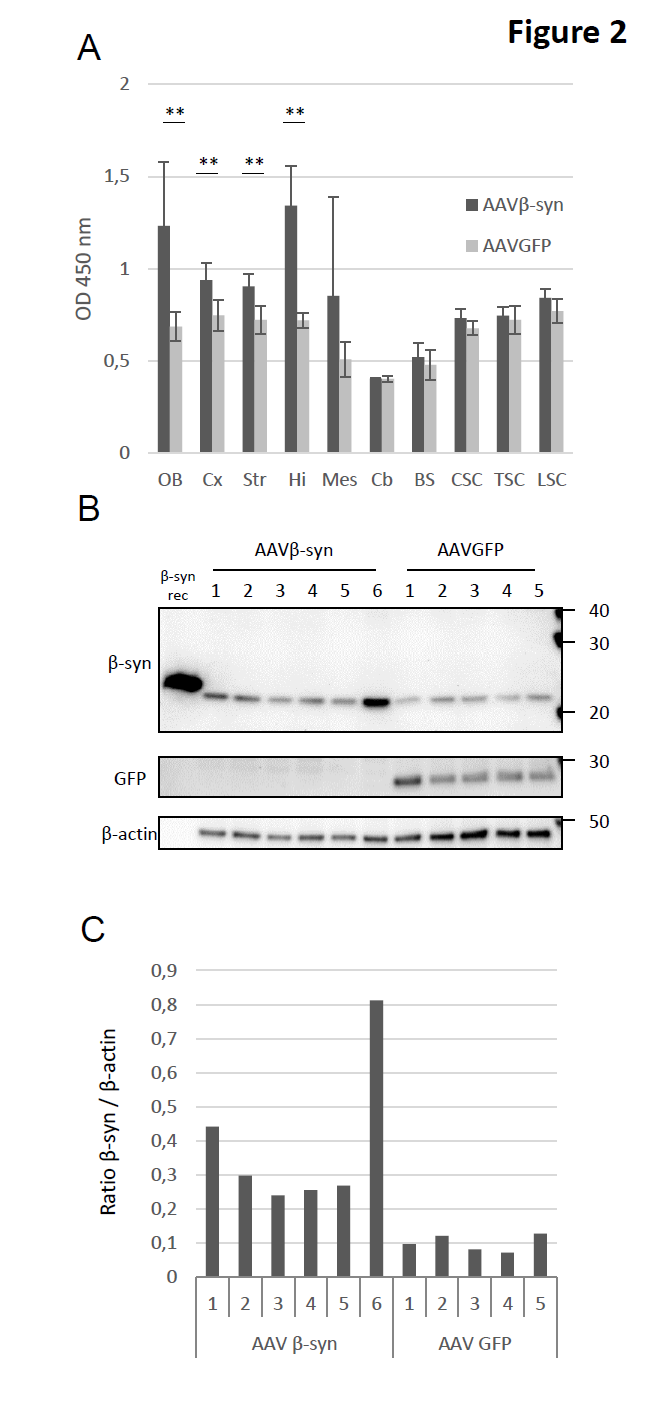
Characterization of the AAV vectors expression after their intracerebroventricular inoculation in M83 mice. M83 mice were injected with AAV vectors in the ventricles at birth and challenged with MSA/M83 inoculum (same mice than in Figure 1 E-G). (A) Detection of total β-syn synthesis by ELISA in cerebral regions and spinal cord of sick M83 mice 5 to 7 months after ICV injection of AAV. (B) Detection of total β-syn or GFP protein in the hippocampus of the same M83 mice by Western blot. Recombinant β- syn was loaded as a positive control. Numbers refer to each mouse (n=6 in the treated group and 5 in the control group). (C) Quantification of total β-syn levels detected by WB. OB: olfactory bulbs, Cx: cerebral cortex, Str: striatum, Hi: hippocampus, Mes: mesencephalon, BS: brain stem, CSC: cervical spinal cord, TSC: thoracic spinal cord, LSC: lumbar spinal cord. Data are shown as means ± sd. *p<0,05; **p<0,01 according to Wilcoxon test.

We next examined the possible effects of AAVβ-syn ICV injection on the spontaneous development of the M83 disease during aging, without any challenge to accelerate the disease (Figure 3A). Here again, AAVβ-syn injection neither delayed the disease onset of M83 mice nor modified the α-syn^P^ levels (Figure 3B and D), despite a sustained expression of the transgene, at least in the hippocampus, up to the disease onset (Figure 3C) (ELISA data correspond to 2 mice for the treated group and 3 mice for the control group; consequently, no statistical analysis was done here).

**Figure 3.**
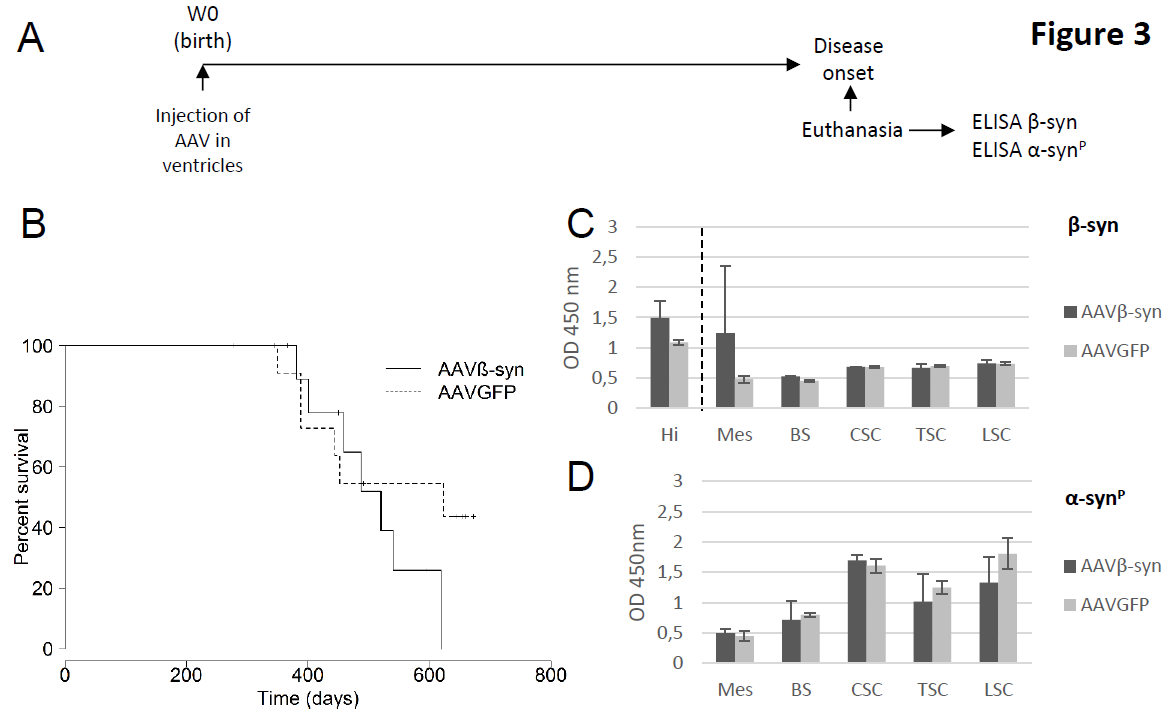
Impact of the intracerebroventricular inoculation of AAVβ-syn or AAVGFP on the spontaneous M83 disease. (A) M83 mice were injected at birth (Week 0) with AAVβ-syn or AAVGFP vectors per lateral ventricle. After the appearance of the first specific symptoms of M83 disease (from 11 months to 20 months of age), mice were sacrificed and dissected in order to perform biochemical analysis. (B) M83 disease-associated survival after the intracerebroventricular (ICV) inoculation of AAV vectors at birth (no significant difference according to log-rank test). (C, D) Quantification of total β-syn (B) or α-syn^P^ (C) by ELISA in CNS regions of the same sick M83 mice. No statistical analysis was done for ELISA results because the number of mice analyzed was too small (2 mice for the treated group and 3 mice for the control group). Hi: hippocampus, Mes: mesencephalon, BS: brain stem, CSC: cervical spinal cord, TSC: thoracic spinal cord, LSC: lumbar spinal cord. Data are shown as means ± sd (n=2 and 3).

Altogether, these results indicated that AAVβ-syn injection in the ventricles resulted in a widespread expression of the transgene in the CNS, but at a moderate level, especially in the structures that are the most heavily affected by the synucleinopathy lesions. We thus considered another strategy by inoculating the AAV vectors in the ventral tegmental area (VTA), which is located into the mesencephalon, where major brain lesions and accumulation of α-syn^P^ are detected in sick M83 mice and which is also connected to multiple brain areas 13.

In order to validate this injection protocol of AAV vectors into the VTA, two months old wild-type mice were first inoculated with AAVβ-syn in the VTA and sacrificed one month later. After dissection of the brains, vector mRNA was mostly detected in the mesencephalon, but also in the cortex, striatum, hippocampus and brainstem; only traces of vector mRNA were identified in the other regions of the brain (except the olfactory bulbs) and in the spinal cord, as compared to mesencephalon (Supplementary figure 1C).

The same protocol was then applied to M83 mice. After the inoculation of AAV in the VTA in two months old M83 mice, the disease was accelerated by intracerebrally inoculating the mice one month later with the M83/M83 or the MSA/M83 inoculum (Figure 4A). No significant difference in the survival was observed between the AAVβ-syn and AAVGFP injected mice, between 3 and 8 months (Figure 4B, E). Importantly, analyses by ELISA confirmed the strong overexpression of β-syn in the mesencephalon although no difference was detected in the levels of pathological α-syn^P^ at the disease stage (Figure 4C, D, F, and G). By immunohistochemistry, using an antibody targeting total β-syn (human and murine β-syn), a specific β-syn staining punctate pattern was detected in the inoculated mesencephalon, as well as in the striatum and brain stem which represent connected regions^13^, of all the sick M83 mice inoculated with AAVβ-syn (5/5), but not in sick M83 mice inoculated with AAVGFP (0/2) (Figure 4H). Further analysis indicated that these β-syn immunoreactive dots, which are specifically detected in mice inoculated with AAVβ-syn, mostly co-localize with the presynaptic protein synaptophysin (Supplementary figure 2A). Co-localization of GFP with neuronal marker β-tubulin type 3 in sick M83 mice inoculated with AAVGFP also confirmed that these vectors allowed a neuronal specific expression of the transgene (Supplementary figure 2B). We further biochemically characterized vectors expression after injection in the VTA in M83 mice euthanized 4 months later. Total β-syn levels were quantified by ELISA and showed a significant increase of the protein in the mesencephalon of mice injected with AAVβ-syn as well as in the hippocampus and brain stem compared to control mice injected with the AAVGFP (Figure 5A). β-syn protein overexpression was also readily detected by Western blot in the mesencephalon of most of these mice (Figure 5B).

**Figure 4.**
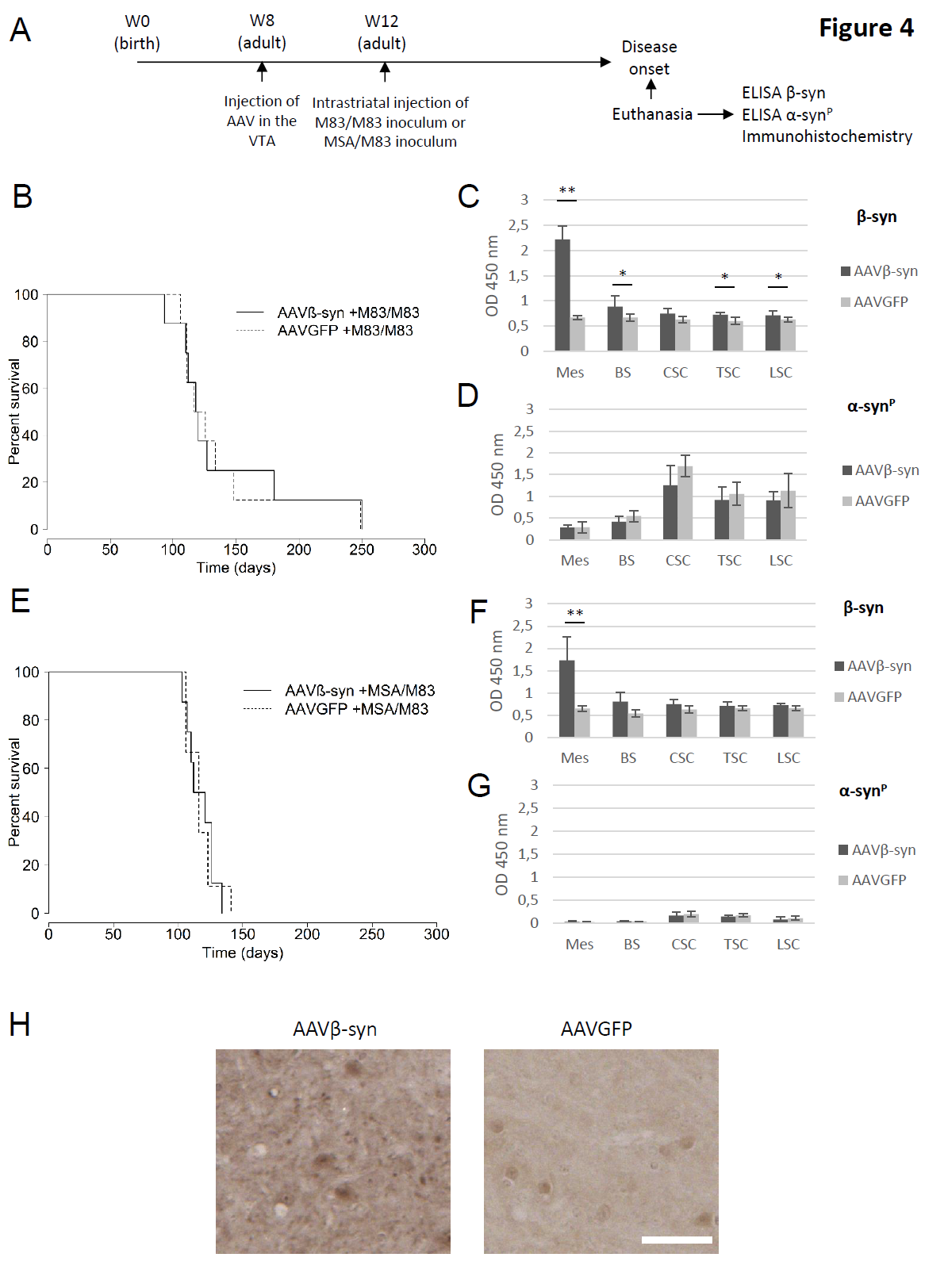
Impact of the inoculation of AAV in the ventral tegmental area on M83 disease. (A) Two months old M83 mice (Week 8) were injected with AAVβ-syn or AAVGFP vectors in the ventral tegmental area (VTA) and challenged one month later (Week 12) by the injection of M83/M83 (B-D) or MSA/M83 (E-G) inoculum in the striatum to accelerate the disease. As before, after the appearance of the first specific symptoms of M83 disease (from 3 months to 8 months after challenge), mice were sacrificed and the CNS was dissected in order to realize biochemical analysis and immunohistochemistry /immunofluorescence studies (Figure 4H, Supplementary figure 2, 3). (B) M83 disease-associated survival after the inoculation of the brain extract (no significant difference according to log-rank test). (C, D) Quantification of total β-syn (C) or pathological Ser129-phosphorylated form of α-syn (α- syn^P^) (D) by ELISA in CNS regions of the same sick M83 mice groups (n=5 for the treated group, n=5 for the control group). (E) The M83 disease-associated survival after the inoculation of the MSA/M83 inoculum (no significant difference according to log-rank test). (F, G) Quantification of total β-syn (F) or α-syn^P^ (G) by ELISA in CNS regions of the same mice groups (n=5 for the treated group, n=5 for the control group). (H) Immunohistochemistry pictures showing total β-syn staining (mouse β-syn and human β-syn produced by the vector) in the mesencephalon of 7 months old sick M83 mice inoculated with AAV β-syn or AAV GFP and challenged with MSA/M83 inoculum. Mes: mesencephalon, BS: brain stem, CSC: cervical spinal cord, TSC: thoracic spinal cord, LSC: lumbar spinal cord. Data are shown as means ± sd. *p<0,05; **p<0,01 according to Wilcoxon test. Scale bar 100μm.

**Figure 5.**
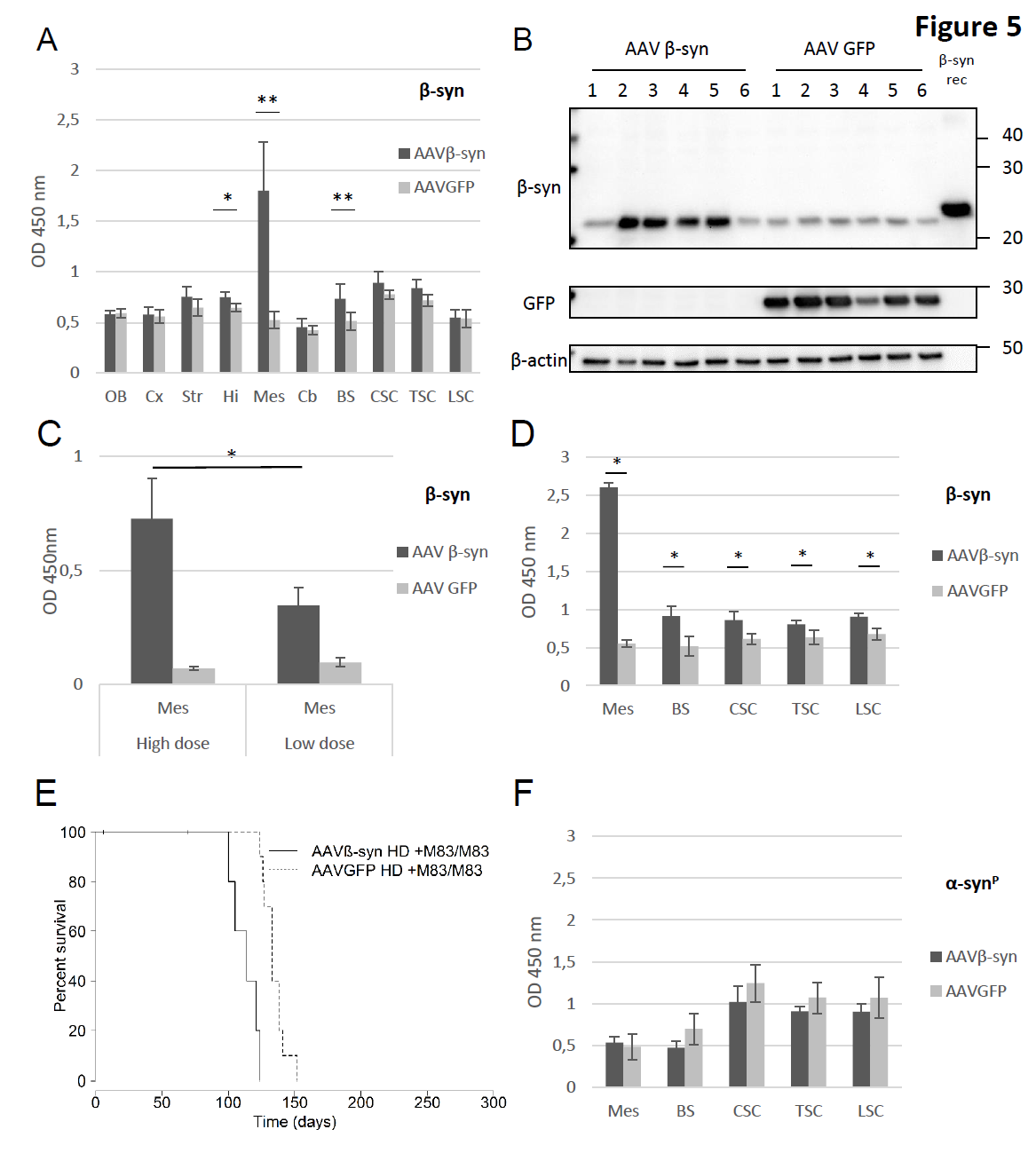
Characterization and impact of inoculation of AAV in the ventral tegmental area of M83 mice. (A-B) Two months old M83 mice were injected with AAVβ-syn or AAVGFP vectors in the VTA and challenged one month later by the injection of M83/M83 inoculum in the striatum to accelerate the disease. In this experiment, all mice were euthanized 3 months after the challenge for biochemical analysis. (A) Detection of β-syn and GFP proteins by ELISA in the brain and the spinal cord of M83 mice after inoculation of AAVβ-syn or AAVGFP in the VTA (n=4 and 6, respectively). (B) Detection of the total β- syn protein by Western blot in the mesencephalon of the same sick M83 mice. Recombinant β-syn was loaded as a positive control. Numbers refer to mice. (C) Total β-syn protein quantification by ELISA test using 0,2 μg (instead of 2 μg) of mesencephalon homogenates extracted from sick M83 mice inoculated with 1,5*10^9^ vg (high dose, 4-times more than the low dose) or 3,75*10^8^ vg (low dose) of AAVβ-syn or AAVGFP. These results were obtained with the same samples used in Figure 4C and 5D, but 10 times diluted. (D-F) Two months old M83 mice (Week 8) were injected with high dose of AAV vectors (4-times higher than in Figure 4 for each vector) in the VTA and challenged one month later (Week 12) by injecting M83/M83 inoculum in the striatum. (D, F) Quantification of total β-syn (D) or α-syn^P^ (F) by ELISA in CNS regions of the same sick M83 mice groups (n=3 and 7 for the treated and control group). (E) M83 disease-associated survival after the inoculation of the brain extract (significant difference according to log-rank test, p<0,001). Mes: mesencephalon, BS: brain stem, CSC: cervical spinal cord, TSC: thoracic spinal cord, LSC: lumbar spinal cord. Data are shown as means ± sd. *p<0,05, **p<0,01, according to Wilcoxon test. Scale bar 100μm.

We finally asked whether the absence of any significant effect of AAVβ-syn could be due to an insufficient dose of AAV vector. We thus injected 4-times more vector in the VTA one month before the challenge by intra-cerebral inoculation of the M83/M83 inoculum. This AAV dose allowed to significantly increase the overexpression of β-syn at the disease stage (p<0,05) (Figure 5C, D). However, the M83 disease was still not delayed by β-syn overexpression, but instead, as previously observed after ICV injection of AAV (Figure 1B), the disease appeared significantly earlier in the AAVβ-syn injected mice as compared to control AAVGFP animals (p<0,001) (Figure 5E). As in previous experiments, no impact was found on the α-syn^P^ levels even if β-syn was significantly increased at the stage of clinical signs in all the CNS regions (Figure 5D, F).

By immunohistochemistry, as with the low dose of AAV, a specific β-syn staining punctate pattern was detected in the inoculated mesencephalon, as well as in the striatum and brain stem in two sick M83 mice inoculated with AAVβ-syn, but not in one mouse inoculated with AAVGFP (data not shown).

Proteinase K (PK)-resistant β-syn aggregates have been previously described after inoculation of AAV vectors expressing human β-syn in the *substantia nigra* of rats^10,11^. We thus analyzed by immunohistochemistry the brain of several sick M83 mice after inoculation of AAVs in the VTA (Figure 6). After PK digestion, total β-syn antibody revealed a punctate pattern specifically detected after the inoculation of AAVβ-syn, in the mesencephalon of 7/7 sick M83 mice (5 mice injected with low dose of AAV, of which 2 were challenged with M83/M83 inoculum and 3 with MSA/M83 inoculum, as well as 2 mice injected with high dose of AAV and challenged with M83/M83 inoculum). This punctate pattern was also detected in the striatum and the brain stem where the AAV is also expressed, but not in the cerebellum where the AAV is less expressed (Supplementary figure 3). A few cell bodies were also stained with total β-syn antibody after PK digestion, particularly in the mesencephalon of these mice (data not shown). Even if diffuse, PK resistant staining was also detected in the hippocampal region of two sick M83 mice injected with AAVβ-syn and not challenged with brain extract inoculation, but not in one sick M83 control mouse (data not shown). These results confirm that overexpression of β-syn resulted in PK-resistant β-syn staining which could be detected independently of the vector dose, the inoculation site or the age of inoculation.

**Figure 6.**
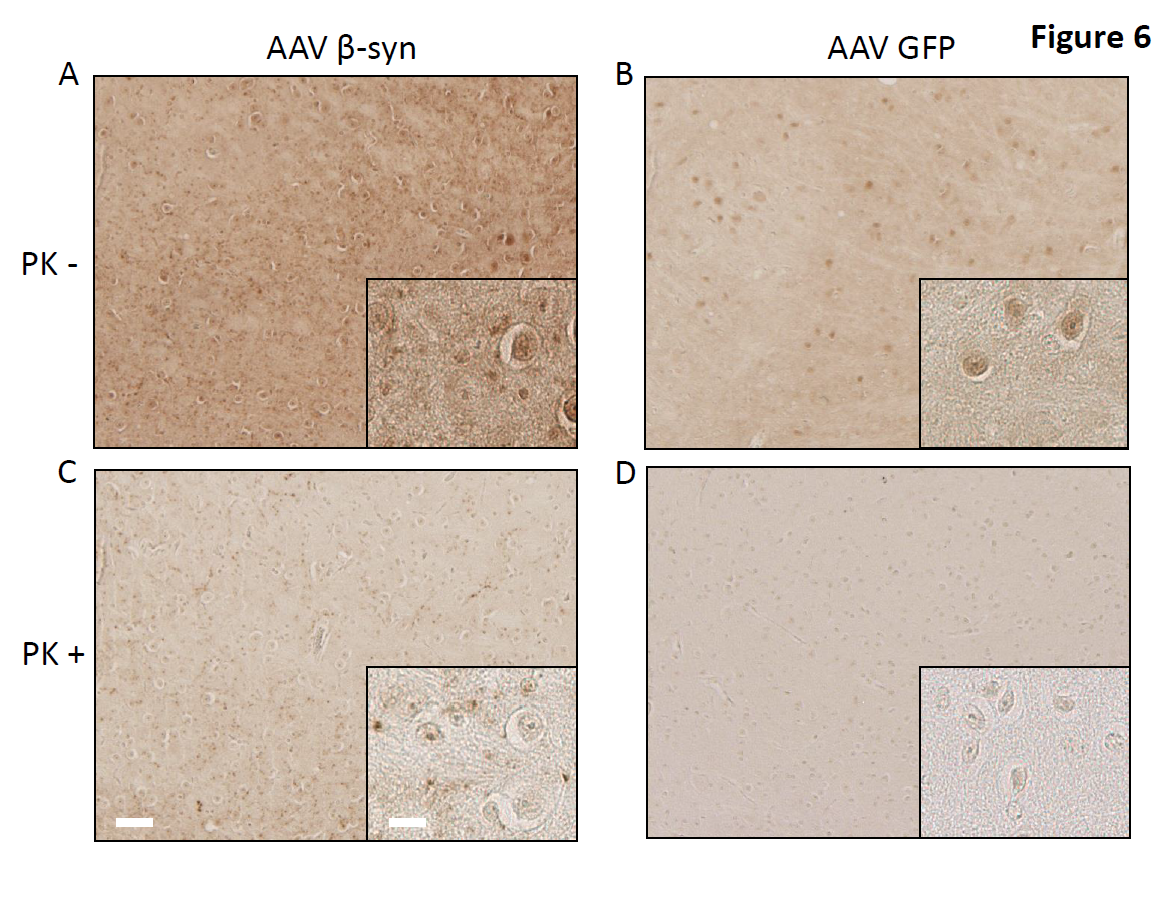
Proteinase K (PK)-resistant β-syn staining in sick M83 mice inoculated with AAV β-syn. (A-D) Immunoreactivity to total β-syn antibody in mesencephalon sections of two sick M83 mice inoculated with low dose of AAVβ-syn or AAVGFP in the VTA and challenged by MSA/M83 inoculum (mice from the study Figure 4E-G) is shown without PK digestion (A, B) or after PK digestion (C, D). After PK digestion, these β-syn immunoreactive dots were detected in all mice analyzed by immunohistochemistry inoculated with the AAVβ- syn in the VTA (7/7 mice, comprising 2 mice challenged with M83/M83 inoculum, 3 mice challenged with MSA/M83 inoculum, and also in 2 mice inoculated with high dose of AAV and challenged with M83/M83 inoculum), but not in sick M83 mice inoculated with AAVGFP (3/3, comprising 2 mice challenged with MSA/M83 inoculum and 1 mouse challenged with M83/M83 inoculum). Scale bars 100μm (low magnification) and 25μm (high magnification).

## Discussion

β-syn was previously described as a neuroprotective protein, able to inhibit the aggregation process of α-syn *in vitro* and having benefic effects on synucleinopathies *in vivo*^6-9,15^. Here, we tried to slow down a synucleinopathy in a transgenic mouse model with dramatic clinical symptoms linked to pathological α-syn aggregation. In particular, we analyzed the effects of the overexpression of β-syn mediated by an AAV vector on α-syn aggregation. We show here that the delivery of the AAVβ-syn vector *via* two different injection routes (intracerebroventricular or in the ventral tegmental area) in transgenic M83 mice, neither delayed the disease onset nor reduced the levels of pathological α-syn at the disease stage, despite a sustained overexpression of β-syn. A previous study suggested that constitutive overexpression of β-syn in transgenic M83 mice by generating double transgenic mice delays the M83 disease onset by several months^9^. In our study, interestingly, M83 mice overexpressing β-syn showed a higher spontaneous activity in experiments in which open field was performed (Supplementary figure 4). These results suggest that β-syn expression might have an impact on mice behavior without affecting the ultimate development of the disease. Importantly, in this study of Fan and colleagues, human β-syn was expressed from the pan-neuronal mouse prion promoter, which is the same promoter used to express human mutated α-syn in the M83 model. It is possible that the strategies tested here did not provide sufficient expression of β-syn, in the CNS regions targeted by the disease. Indeed, as expected, after the injection of AAV in the VTA, we obtained a very high overexpression of β-syn in the mesencephalon in which the aggregation of α-syn is high at the disease stage, but this region may not be the region where the aggregation process of α-syn begins^16^. It should be emphasized that the major signs of the clinical disease of M83 mice are in relation with some - still poorly explained-disorders in the spinal cord^17^. In an additional series of experiments, we were unable to identify a degeneration of motor neurons in the lumbar spinal cord after the injection of brain extract from sick M83 mouse or preformed fibrils of human α- syn in M83 mice, as previously reported after intra-muscular injections^17^, further interrogating the cause of the appearance of symptoms (Supplementary figure 5). The ICV injection of AAV9 vector in SMNΔ7 mice, a severe model of spinal muscular atrophy, allowed to express a codon-optimized human SMN1 coding sequence in the spinal cord, which as a result, has improved significantly their survival^18^. After the ICV inoculation of AAV β-syn, the overexpression of β-syn was diffuse and lower in caudal regions of the brain and spinal cord, as described^14^, but it may have been insufficient to slow down the synucleinopathy process. It must however be noted that an *in vitro* study using AAV vectors suggests that β-syn overexpression is benefic on α-syn linked toxicity only at a low ratio of β- syn/α-syn expression^10^. Even if it was not evaluated here, another study has described transgene expression in the neuromuscular junction after AAV injection into the ventricles at birth; this may be important since the degeneration of the neuromuscular junction was also described as a possible explanation of the appearance of symptoms in M83 mice^14,17^. Another study suggested that the benefic effect of β-syn is the result of the down regulation of the expression of α-syn, but we did not detect any decrease of α-syn expression, even in the brain regions overexpressing β-syn (data not shown). In another transgenic mouse model (D line) overexpressing human wild type α-syn, two studies reported the reduction of the synucleinopathy by either generating bigenic mice that also overexpress β-syn, or vehiculating the human β-syn transgene, under the control of cytomegalovirus (CMV) promoter, in a lentiviral vector^7,8^. As some reports indicate a downregulation of the CMV promoter in the CNS over time^19^, we chose here to use the human synapsin 1 promoter to obtain a long term expression of β-syn, which was confirmed in the present study, even in old mice. Importantly, as previously described^20^, the synapsin 1 promoter also allowed to express human β-syn specifically in neurons, where α-syn aggregates are found in the M83 model^1^. According to the study by Hashimoto *et al.*, the neuroprotective effect of β-syn was associated to the activation of Akt, a neuroprotective protein which could be indirectly down-regulated by α-syn during the synucleinopathy process^21^. However, we did not detect any significant increase of Akt activation by Western blot in 6 months-old M83 mice inoculated with AAVβ- syn in the VTA at 2 months, and challenged one month later with the M83/M83 inoculum (Supplementary figure 6). It should be noticed that such Akt changes were not consistently observed, as shown in a study with a lentiviral vector used for β-syn overexpression^22^.

Surprisingly, in the present study, two experiments in mice challenged with a M83/M83 inoculum rather showed a significant acceleration of the disease after AAV β-syn injection (Fig 1B and 5E). Interestingly, several studies reported that β-syn could be implicated in the pathological process of synucleinopathies. Indeed, an axonal pathology with β-syn accumulation in axons was reported in patients with Parkinson’s disease, DLB or Hallervorden-Spatz syndrome^23,24^. In addition, two mutations of β-syn, identified in DLB cases, were suggested to be responsible for α-syn aggregation and DLB^25^. Generating transgenic mice co-expressing P123H mutated β-syn with human α-syn resulted in an enhanced pathology, suggesting that mutated β-syn could potentiate α-syn aggregation^26^. Variable results were also observed after overexpression of β-syn using a lentiral vector after inoculation into the hippocampus of transgenic mice overexpressing a mutated form of the human β-amyloid precursor protein (APP) involved in Alzheimer’s disease, as assessed by examining plaque load, memory deficits, and anxiety^22^. However, it should be noted that the expression level of β-syn was relatively low in this latter study (not detectable at the protein level). Also, it is interesting to notice that a truncated form of human recombinant α-syn lacking the 71-82 residues not able to form aggregates *in vitro*, like β-syn^27^, accelerated the disease of a few M83 mice, after injection in the muscle or in the peritoneal cavity^17,28^. These results raise the possibility that β-syn might be able to accelerate the aggregation of α-syn in M83 mice, although this has not been reported so far.

Further analyzing the effects of β-syn expression in sick M83 mice inoculated with the AAV vector in the VTA allowed detecting a punctate pattern of β-syn staining in the targeted brain region and some connected regions, only in mice inoculated with AAVβ-syn. This staining was more resistant to PK digestion than that endogenous β-syn in control mice. This result suggests that overexpressed β-syn formed aggregates which could explain why β-syn did not act as expected, *i.e* as an α-syn aggregation inhibitor. Landeck *et al.* have described the same staining of β-syn resistant to PK digestion referred as «dark punctae«, two months after the inoculation of AAV5 carrying human β-syn under the control of chicken beta-actin promoter including a CMV enhancer element (CBA) in the *substantia nigra* of rats^11^. Before this study, Taschenberger *et al.* have also described the detection of PK-resistant β-syn aggregates *in vivo*, as early as two weeks following the inoculation of AAV2 carrying human β-syn gene in the *substantia nigra* of rats^10^. This time, the vector design was closer to ours, using the same promoter (human synapsin) to express human β-syn, with a WPRE sequence to enhance transgene expression. It is interesting to notice that, even if the AAV serotype used as well as the vector design can differ between studies, all these studies that used AAV to carry β-syn also reported the detection of β-syn aggregates. These observations suggest that the neuroprotective effect of β-syn may vary according to the strategy followed to overexpress it and the animal model. Also, it is important to point out that even if most of neurons express the transgene in the cerebral region injected, there is an important variability of the expression of the transgene in transduced cells after AAV vectors injection (Supplementary figure 7). It might explain at least partly the difficulty to detect a benefic effect of β-syn overexpression as it was suggested to depend on a specific α-syn/β-syn ratio in cell culture^10^.

The present study suggest for the first time that β-syn overexpression could not be benefic in a synucleinopathy model. This may be due at least partly to the formation of a PK-resistant β- syn species after AAVβ-syn inoculation. According to a very recent study, a mildly acidic pH environment, found in several organelles, could induce β-syn aggregation, highlighting the complexity of β-syn fibrillation mechanisms occurring *in vivo*^29^. Further studies are needed to better understand how and to what extent β-syn could play a role in synucleinopathies.

However, even though we could not detect any protective effects of β-syn, our study confirms that an AAV vector is suitable for long-term overexpression of proteins into the CNS. Notably, after ICV inoculations of AAV in neonates, we confirmed a widespread expression of the protein, as this was also recently described using an α-syn AAV vector^30^. The injection of AAV in the VTA also allowed to express human β-syn in various connected regions, as suggested in a previous study using serotype 9 AAV vector carrying a lysosomal enzyme gene^13^. Our data also illustrate the robustness of the experimental model of M83 mice intracerebrally challenged by inocula containing aggregated α-syn, showing rather short and relatively uniform survival periods before the appearance of characteristics clinical signs, which can be used for the assessment of future therapeutic strategies.

## Methods

### Animals

M83 transgenic mice were used in this study (B6;C3H-Tg[SNCA]83Vle/J, RRID:MGI:3603036, The Jackson Laboratory, Bar Harbor, ME, USA). These mice express A53T mutated human α-syn protein and spontaneously develop severe motor impairment leading to early death^1^. Homozygous M83 mice develop characteristic motor symptoms between 8 and 16 months of life, beginning with reduced ambulation, balance disorders, partial paralysis of a hind leg and then progressing to prostration, difficulty in feeding, weight loss, hunched back and general paralysis^1^. The animals were housed per group in enriched cages in a temperature-controlled room on a 12h light/dark cycle, and received water and food *ad libitum,* in our approved facilities (No. C69 387 0801) for breeding and experimental studies, in accordance with EEC Directive 86/609/EEC and French decree No. 2013-118. The experimental studies described in this article were performed in containment level 3 facilities and authorized by the « Comité d’éthique » CE2A – 16, ComEth ANSES/ENVA/UPEC and by the « Ministère de l’enseignement supérieur, de la recherche et de l’innovation » (ref 16- 006).

### AAV vectors

Recombinant self-complementary AAV9 vectors (scAAV9) encoding human β- syn or enhanced GFP (eGFP) were produced by calcium phosphate transfection of HEK-293 cells^31^. Three plasmids were transfected simultaneously: (i) a vector plasmid containing the human gene of β-syn or eGFP under control of the neuron-specific synapsin 1 gene promoter (Figure 1A) (ii) a helper plasmid pXX6^32^ and (iii) a plasmid carrying rep2 and cap9 genes^33^. Vector particles were extracted and purified on an iodixanol step gradient. Titers were determined by quantitative polymerase chain reaction (qPCR) and expressed as viral genomes per milliliter (vg/mL).

### Inoculations

AAV vectors were injected at birth or at adulthood (two months of age), at different sites of inoculation. Neonates were inoculated using 5 μL Hamilton syringe, with 9.38*10^8^ vg of AAVβ-syn or 2.73*10^8^ vg of AAVGFP vectors per lateral ventricle of the viral solution with 0.05% Trypan blue^12^. Adults mice were inoculated with 3.75*10^8^ vg of AAVβ-syn or 1.09*10^8^ vg of AAVGFP (or with high dose: 1.5*10^9^ vg of AAVβ-syn or 4.36*10^8^ vg of AAVGFP) of viral solution in the ventral tegmental area located in the mesencephalon, using stereotaxic coordinates (anteroposterior: −3.16 mediolateral: +0.25 dorsoventral: −4.50). Two months after the inoculation of the viral solution in neonates or one month after the inoculation of the viral solution in adults, M83 mice were inoculated with brain homogenates from a second passage of a sick M83 brain sample in M83 mice (M83/M83 inoculum) or a second passage of a patient with multiple system atrophy (MSA) brain sample in M83 mice (MSA/M83 inoculum) (same sample used in a previous publication 5). These homogenates were injected in the left striatum, using stereotaxic coordinates (AP: +0.14 ML: +2 DV: −2.75). Before each stereotaxic surgery (AAV or brain extract inoculation), mice have been anesthetized with a xylazine (10 mg/kg) and ketamine (100 mg/kg) mixture.

### qRT-PCR

B6C3H brains were dissected and RNA was extracted from brain regions using RNeasy lipid tissue Minikit (ref 74804, Qiagen, Courtaboeuf, France). RNA samples were tittered using Nanodrop (Nanodrop 1000, Thermo Fisher scientific, Villebon sur Yvette, France). 500 ng of RNA were reverse-transcripted with Quanta qScript cDNA Super Mix (ref 95048-100, VWR international, Fontenay-sous-Bois, France) on a Biorad iCycler. After a 1:10 dilution, complementary DNA samples were analyzed by qPCR using primers specific for the WPRE region of viral mRNA, and primers specific for GAPDH mRNA. We used the LC 480 SYBR Green 1 Master kit (ref 04887 352 001, Roche, Boulogne-Billancourt, France) on a Roche Light Cycler 480 to perform the qPCR and analyzed the results on the LC 480 software.

### ELISA

Brains were dissected as described^4^ and proteins were extracted in high salt buffer (50 mM Tris-HCl, pH 7.5, 750 mM NaCl, 5 mM EDTA, 1 mM DTT, 1% phosphatase and protease inhibitor cocktails), using a mechanical homogenizer (grinding balls, Precellys 24, Bertin Technologies, Montigny-le-Bretonneux, France) to obtain a 10% homogenate (w/v). Each sample was titered using the DC^TM^ protein assay kit (ref 5000111, Biorad, Marnes-la-Coquette, France).

As already described for α-syn^P^ detection^3,5^, plates were saturated with Superblock T20 (Thermo Scientific, Rockford, IL, USA) for 1 h at 25°C, under agitation (150 rpm). After 5 washes in PBST, 10 μg of protein (for α-syn^P^ detection) diluted in PBST were incubated for 2 h at 25°C, under agitation (150 rpm). α-syn^P^ was detected with a rabbit polyclonal antibody against PSer129 α-syn (ref ab59264, Cambridge, UK) diluted to 1:3,000 in PBST with 1% bovine serum albumin (BSA); plates were incubated for 1 h at 25°C under agitation. After 5 washes, anti-rabbit IgG HRP conjugate (ref 4010-05, SouthernBiotech, Birmingham, AL, USA) was added at 1:2,000 (for α-syn^P^ detection). After washing, 100 μL of 3,3′,5,5′- tetramethylbenzidine solution (ref. T0440, Sigma, Saint-Quentin-Fallavier, France) were added to each well and plates were incubated for 15 min with shaking. The reaction was stopped with 100 μL of 1 N HCl, and the absorbance was measured at 450 nm with the microplate reader Model 680 (Clariostar, BMG Labtech, Champigny sur Marne, France).

ELISA allowing total β-syn quantification was adapted based on an ELISA already published 22. Briefly, 2 μg or 0.2 μg (for Supplementary figure 3) of brain homogenate was diluted in PBST BSA 1% and incubated at 4°C overnight. After 5 washes in PBST, plates were saturated with Superblock T20 for 1 h at 25°C, under agitation (150 rpm). After 5 washes in PBST, plates were incubated with 1:2,000 of an anti-β-syn antibody ab76111 (ref EP1537Y, Abcam, Cambridge, UK). After 5 washes in PBST, plates were incubated with an anti-rabbit IgG HRP conjugate (ref 4010-05, SouthernBiotech, Birmingham, AL, USA) at 1:4,000. After washing, immunoreactivity was revealed with the same protocol as for α-syn ELISA.

### Western blot

50 μg (for phosphorylated Akt detection) or 10 μg of proteins (for total Akt detection) or 2 μg of proteins (for GFP and β-syn detection) were separated in 12% SDS-polyacrylamide gels and electroblotted onto polyvinylidene fluoride (PvF) membranes 0,45 μm (Bio-Rad). The membranes were washed 3 times in Tris-Buffered Saline (TBS) for 5 min at room temperature (RT) under agitation and were saturated 1 h with 5% BSA in TBS 0.1% Tween20 (TBST). Membranes were incubated with rabbit antibody against phosphorylated Akt at Ser473 (ref 9271S, Ozyme, Montigny-le-Bretonneux, France) at 1:1,000 or with rabbit antibody against total Akt (ref 9272, Ozyme, Montigny-le-Bretonneux, France) at 1:1,000 or with anti-β-syn antibody ab76111 (ref EP1537Y, Abcam, Cambridge, UK) at 1:5,000 or with anti-GFP antibody (ref ab290, Abcam, Cambridge, UK) at 1:1,000 or with anti-β-actin antibody (ref ab8226, Abcam, Cambridge, UK) at 1:2000 overnight at 4°C. After 3 washes, the membranes were incubated for 1 h at RT with anti-rabbit HRP-linked antibody (ref 7074P2, Ozyme, Montigny-le-Bretonneux, France) for detection of total Akt and phosphorylated Akt at 1:2,000, or with anti-rabbit HRP-linked antibody (ref 4010-05, SouthernBiotech, Birmingham, AL, USA). The immunocomplexes were revealed with chemiluminescent reagents (Supersignal WestDura, ref 34076, Pierce, Interchim, MontLuçon, France), and analyzed using the ChemiDoc system (Bio-Rad) and Image Lab software (Bio-Rad) which allowed to quantify the intensity of the bands.

### Immunohistochemistry/Immunofluorescence

After dissection, brain samples were fixed in 4% paraformaldehyde and paraffin embedded to be cut into serial 6 μm sections. After deparaffinization, endogeneous peroxidase activity was directly blocked with oxygenated water 3% during 5 minutes at RT. For immunofluorescence, brain sections were pretreated with a citrate solution (ref C9999, Sigma, Saint-Quentin-Fallavier, France) with heat antigen retrieval. After washing, sections were saturated using a blocking reagent (ref 11096176001, Roche), 1 h at RT and incubated with antibody against β-syn ab76111 in PBST (ref EP1537Y, Abcam, Cambridge, UK) diluted at 1:500, anti-synaptophysin antibody SY38 (ref ab8049, Abcam, Cambridge, UK) diluted at 1:10 or anti-GFP antibody (ref ab290, Abcam, Cambridge, UK) diluted at 1:500 with anti-tubulin β-3 antibody (ref MMS-435P, Biolegend, San diego, USA) diluted at 1:100 at 4°C overnight. After another blocking step 30 minutes at RT, sections were incubated 1 h at RT with anti-rabbit IgG HRP conjugate (ref 4010-05, SouthernBiotech, Birmingham, AL, USA) diluted at 1:250 in PBST. Antibody binding was detected using DAB peroxidase substrate (ref SK-4100, Vector Laboratories, Burlingame, CA USA) intensified with nickel chloride. For immunofluorescence, sections were incubated 1 h at RT with anti-rabbit IgG AlexaFluor 488 or 555 (ref A-11008 and ref A21428, respectively, Thermofischer scientific, Villebon sur Yvette, France) or with anti-mouse IgG AlexaFluor 488 or 555 (ref A11001 and A21127 respectively, Thermofischer scientific, Villebon sur Yvette, France) diluted at 1:1000. After washings in PBST then in PBS, sections were treated with an autofluorescence eliminator reagent (ref 2160, Millipore, Temecula, USA), before mounting.

### PK digestion

After being deparaffinized, sections were incubated with a 10 μg/mL solution of proteinase K (ref EU0090-B, Euromedex, Souffelweyersheim, France) diluted in PBS, 10 minutes at RT (protocol adapted from^10^). Sections were washed three times in water before blocking the endogenous peroxidase activity and pursuing the immunohistochemistry staining protocol.

### Statistical analysis

Survival time was defined as the time from birth until death (Figure 3A, exclusively) or as the time from the inoculation of the brain extract until the appearance of the first specific M83 symptoms and euthanasia of the mouse. We right-censored mice found dead without M83 disease identification. Survival times were compared using log-rank test. Concerning statistical analysis of ELISA tests results, means were compared using Student test, when at least 6 animals were included in all groups of animals which were compared or using Wilcoxon test, when at least one group of mice was composed of less than 6 animals. The difference was significant when p<0,05 (*), p<0,01(**), p<0,001(***).

## Acknowledgements

We are grateful to Eric Morignat and Habiba Tlili for their help and advices on statistical and immunofluorescence assays, respectively. We also are grateful for Olivier Biondi for its help setting up behavioral test and the method for quantifying motor neurons in the spinal cord. We thank Ronald Melki and its team for providing us preformed fibrils of recombinant α-syn. This research was partly funded by the Fondation France Parkinson. D.S was supported by funds from the Région Auvergne-Rhône-Alpes – ARC1 Santé.

## Author contributions

DS, DB, MD, DG, performed experiments. JV, JNA contributed to biochemical and immunohistochemistry assays respectively. LL supervised animal experiments in containment level 3 facilities. AS supervised all the construction and the production of viral vectors. TB, DB designed experiments and supervised the study. DS, TB, DB, AS wrote the paper. All authors read and approved the final manuscript.

